# Exploring the mechanistic link between corticosterone and insulinlike growth factor-1 in a wild passerine bird

**DOI:** 10.1101/306407

**Authors:** Zsófia Tóth, Jenny Q. Ouyang, Ádám Z. Lendvai

## Abstract

**Background:** Physiological regulators of life history trade-offs need to be responsive to sudden changes of resource availability. When homeostasis is challenged by unpredictable stressors, vertebrates respond through a set of physiological reactions, which can promote organismal survival. Glucocorticoids have been traditionally recognized as one of the main regulators of the physiological stress response, but the role of an evolutionarily more conserved pathway, the hypothalamic-pituitary-somatotropic (HPS) axis producing insulin-like growth factor-1 (IGF-1) has received much less attention. Although IGF-1 is known to affect several life history traits, little is known about its role in the physiological stress response and it has never been studied directly in adult wild animals.

**Methods:** In this study, we combined field observations with a controlled experiment to investigate how circulating levels of IGF-1 change in response to stress and whether this change is due to concomitant change in glucocorticoids in a free-living songbird, the bearded reedling *Panurus biarmicus*. We used a standard capture-restraint protocol in field observation, in which we took first and second (stress induced: 15 minutes later) samples. In a follow-up experiment, we used a minimally invasive oral corticosterone manipulation.

**Results:** We showed that corticosterone levels significantly increased while IGF-1 levels significantly decreased during capture and handling stress. However, change in corticosterone levels were not related to change in IGF-1 levels. We found that experimentally elevated corticosterone levels did not affect IGF-1 levels.

**Discussion:** Our results are the first to highlight that circulating IGF-1 levels are responsive to stress independently from glucocorticoids and suggest that the HPS axis is an autonomous physiological pathway that may play an important role as regulator of life-history decisions.

## Introduction

Resource allocation trade-offs are central to the evolution of life-histories. Physiological mediators of such trade-offs need to monitor resource availability and transmit a signal to relevant parts of the organism to adjust energy expenditure in face of environmental variation. Possible candidates of such key life-history regulatory mechanisms must therefore integrate information from both the external and internal environment and be responsive to changes in resource availability (Harshman & Zera, 2007). One way to investigate whether a given physiological mechanism could have such life-history regulatory functions is to assess whether it fulfils these requirements of information processing, integration and responsiveness. A useful framework to analyse these questions is when individuals are exposed to stressors. This approach is biologically and ecologically relevant because in order to successfully reproduce and survive, all organisms must be able to cope with environmental challenges. An organism can display a stress response when challenged by unpredictable, noxious changes in the environment, such as the attack of a predator, an infection, inclement weather, or food shortage (Wingfield et al., 1998; Kitaysky, Wingfield & Piatt, 1999; Romero, Reed & Wingfield, 2000; Hawlena & Schmitz, 2010). Mounting an appropriate stress response requires a dramatic reorganization of resource allocation. Understanding this response provides great insight into life-history decisions and resource allocation mechanisms.

An important regulator of the stress response is the endocrine system, which plays a central role in regulating the adjustment of morphology, physiology and behaviour to deal with current conditions (Ricklefs & Wikelski, 2002; Flatt & Heyland, 2011; Wingfield & Boonstra, 2013). Hormones function as integrators that process information from the environment and orchestrate multiple processes simultaneously to maximise survival and reproduction under the given circumstances (Wingfield et al., 1998; Martin et al., 2011). Although some pathways have been clearly represented in the study of the stress response (e.g. HPA axis - hypothalamic-pituitary-adrenal axis), other, more evolutionarily conserved pathways are understudied even though they can play a key role in the evolution of the endocrine system.

The HPA cascade begins at the brain that perceives the stressor and launches a series of downstream hormonal changes. At the endpoint of this axis, the adrenal cortex secrete increased amounts of glucocorticoids into the bloodstream, which stimulate energy production via gluconeogenesis. However, this pathway appeared during the evolution of vertebrates, by which period robust regulation of homeostasis and resource allocation had already evolved (Stoks, 2001; Baker Michael E., 2003; Bijlsma & Loeschcke, 2005; Pauwels, Stoks & De Meester, 2005; Hawlena & Schmitz, 2010). A more conserved endocrine pathway is the insulin/insulin-like signalling pathway (the IIS pathway), which is present in all animals and regulates resource allocation and stress resistance (Broughton et al., 2005; Harshman & Zera, 2007; Dantzer & Swanson, 2012). In vertebrates, the IIS pathway is integrated into the Hypothalamic-Pituitary-Somatotropic (HPS) axis. As part of this hormonal cascade, growth hormone (GH) stimulates the secretion of insulin-like growth factor-1 (IGF-1), primarily from the liver (Roith, Scavo & Butler, 2001).

IGF-1 is an evolutionarily highly conserved nutrient sensing hormone that has pleiotropic effects influencing key life-history traits and major life-history trade-offs among growth, reproduction and lifespan (Dantzer & Swanson, 2012). IGF-1 is negatively related to lifespan (Holzenberger et al., 2003; Lewin et al., 2016), but has a positive effect on growth, sexual maturation and reproduction (Crain et al., 1995; Yakar et al., 1999; Pine et al., 2006; Flatt et al., 2008; Sparkman, Vleck & Bronikowski, 2009; Lewin et al., 2016, Lodjak, Mägi & Tilgar, 2014, Swanson & Dantzer, 2014). IGF-1 therefore may mediate the trade-off between longevity and reproduction because reduced IGF-1 signalling has been shown to increase lifespan and the expression of genes involved in stress resistance while increased IGF-1 signalling is necessary for reproduction (Holzenberger et al., 2003; Harshman & Zera, 2007; Dantzer & Swanson, 2012; Lewin et al., 2016).

Although the role of IGF-1 has been established in resource allocation, its role in coping with stressful situations and the crosstalk between the HPA and the HPS axes remain poorly understood. It has been shown that IGF-1 levels change under nutritional and handling stress. For instance, in response to 5 minutes of restraint, circulating IGF-1 levels decreased by 21% within 60 minutes in Yorkshire pigs *Sus crofa domesticus* and remained suppressed for up to 150 minutes (Farmer et al., 1991). A more recent study in pigs also found that restraint stress caused a drop in circulating IGF-1 levels and affected other components of the IIS pathway, suggesting that the IGF-system represents a physiologically relevant biomarker of stress response (Wirthgen et al., 2017). Similarly, IGF-1 levels significantly decreased due to short-term (15 min) and long term (24 hours) confinement stress in sunshine bass (hybrid of *Morone saxatilis* and *M. chrysops*), in Atlantic salmon *Salmo salar* and in rainbow trout *Oncorhynchus mykiss* (Wilkinson et al., 2006; Davis & Peterson, 2006).

Whether the decrease in IGF-1 levels is a direct consequence of exposure to stressors on the HPS axis or is due to the stress-induced activity of the HPA axis is controversial (Unterman et al., 1993; Dell et al., 1999; Davis & Peterson, 2006). On the one hand, it is well established that glucocorticoids initiate and orchestrate the emergency life-history stage within minutes to hours from the appearance of the stressor in temperate bird species (Breuner, Greenberg & Wingfield, 1998; Wingfield et al., 1998; Romero & Remage-Healey, 2000; Löhmus, Sundström & Moore, 2006), and we also know that glucocorticoids are interconnected with the somatotropic axis (Dell et al., 1999). For instance, glucocorticoids play a role in the embryonic development of the HPS axis by affecting GH gene expression, IGF-1 transcription (Dell et al., 1999; Bossis & Porter, 2003; Reindl & Sheridan, 2012). Experimental studies have also shown that exogenous glucocorticoids cause lower IGF-1 levels in rats (Gayan-Ramirez et al., 1999), chicken (Leili & Scanes, 1998) and fish (Kajimura et al., 2003; Peterson & Small, 2005). On the other hand, these effects seem to operate only at high glucocorticoid concentrations and/or at a prolonged exposure. For example, although Bossis and Porter (2003) found that glucocorticoids mediate the embryonic development of the HPS axis, hormone treatment for at least 8 hours was necessary to detect a significant increase in GH gene expression. Similarly, while a pharmacological increase in cortisol resulted in a decrease of IGF-1 levels in tilapia *Oreochromis mossambicus* 24-48h post injection (Kajimura et al., 2003), a more moderate, short-term elevation in cortisol did not affect IGF-1 levels in sunshine bass (Davis & Peterson, 2006).

While some of these laboratory, aquacultural and agricultural studies suggest a relationship between stress, glucocorticoids and IGF-1, our knowledge remains very limited about how stressors affect IGF-1 levels in free-living organisms. Importantly, we do not know how the HPA and HPS axes are linked mechanistically. Birds are particularly interesting model systems for studying the IIS, because their metabolism is faster yet their lifespan is longer than similar-sized mammals (Costantini, 2008), and differences in how the IIS is regulated compared to the most studied mammalian models might provide a deeper understanding of the evolution of this pathway (Holmes & Ottinger, 2003; Dantzer & Swanson, 2012). Although the IGF-system has been extensively studied in poultry (reviewed in McMurtry, 1998) almost nothing is known about its regulation in free-living birds (notable exceptions are Lodjak, Mägi & Tilgar, 2014; Lodjak, Tilgar & Mägi, 2016; Lodjak et al., 2017)

While interest in the ecological and evolutionary relevance of IGF-1 has recently increased (Sparkman, Vleck & Bronikowski, 2009; Sparkman et al., 2010; Palacios, Sparkman & Bronikowski, 2012; Lodjak, Mägi & Tilgar, 2014; Reding et al., 2016; Lodjak, Tilgar & Mägi, 2016; Lodjak et al., 2017), to the best of our knowledge no study has directly tested the effects of stressors on the activity of the HPS axis or investigated the mechanistic link between the HPA and HPS axes in any free-living organisms. Therefore, we aimed at answering whether: (1) acute stress affects plasma levels of IGF-1 and whether (2) glucocorticoids directly affect circulating IGF-1 levels. First, we hypothesized that acute stress will affect the activity of the HPS axis; therefore, we predicted that circulating IGF-1 levels will decrease in response to short-term acute stress. Second, to investigate the crosstalk between the HPA and the HPS axes, we experimentally increased corticosterone levels using a minimally invasive technique. We predicted that if the HPS axis receives direct input from glucocorticoids, then increased glucocorticoid levels will down-regulate IGF-1 levels. Alternatively, if the HPS axis responds to external stressors directly, then we predicted that short-term elevation of circulating glucocorticoids will not affect IGF-1 levels. To answer these questions, we studied free-living bearded reedlings *Panurus biarmicus*, a small (~14g), sexually dimorphic, resident songbird common throughout wetlands of Eurasia.

## Material and methods

### 1. Study animal and field study

We captured 17 wintering free-living bearded reedlings *Panurus biarmicus* (Linnaeus, 1758) in Hungary at Virágoskúti-halastó (N 47.6518, E 21.3589) between September 2015 and January 2016. We caught the birds with continuously observed mist-nets and subjected them to a standardized capture-handling-restraint protocol (Wingfield, 1994). The first blood sample (50~100 μl) was taken as soon as possible after the bird hit the net (mean handling time: 4:58 minutes; range: 2-10 minutes). The initial handling times contained one high outlier, inclusion or removal of this point did not affect our results qualitatively. The handling time was not detectably related to either corticosterone levels (t = 1.276, p = 0.223) or IGF-1 levels (t = −0.785 p = 0.445). The bird was then placed into an opaque cloth bag and the collection of a subsequent blood sample (50-100 μl) was started 15 minutes after the initial capture (completed mean time: 17:51 minutes; range: 15-19 minutes). We chose to take the second blood sample after 15 minutes because we were interested in the short-term acute stress-response, and we wanted to minimize the possible downstream effects of corticosterone on IGF-1 levels. The total volume of the two blood samples was under 140 μl that met the recommendation of Owen (2011). The study was approved by the Institutional Animal Care and Use Committee at the University of Debrecen (DEMAB/19-6/2015) and the regional government agency (HBB/17/00870-3/2015).

### 2. Experimental study

#### Housing protocol

Twenty-one wintering bearded reedlings were captured with mist-nets in Hungary at Hortobágy-Halastó (N 47.6211, E 21.0757) between 18 of October and 16 of November in 2016. After capture, they were housed in an outdoor aviary at the Botanical Garden of the University of Debrecen where they were kept for 4 months and acclimated to captivity. Four weeks prior to the experiment, the birds were transferred to individual cages measuring 25×25×25 cm. All sides of the cages (except for the front) were made of a non-transparent board (OSB); therefore, birds were kept in visual but not acoustic isolation. Individual cages were separated by a non-transparent removable divider. Because bearded reedlings are very social, live in flocks and maintain strong pair bonds throughout the year (Lovász, Fenyvesi & Gyurácz, 2017; Griggio & Hoi, 2011), long-term individual separation may be perceived as stressful for the birds (as found in other social species, e.g., Remage-Healey, Adkins-Regan & Romero, 2003). To avoid such additional stressors, before the experiment, birds were kept in pairs, by removing the divider between adjacent cages. Food (a mixture of apple, carrot, quark, cracked dried fish, dried *Gammarus* sp., cracked corn, cracked dry cat food, a commercial soft food mixture for birds and mealworms) and water were available *ad libitum* at all times.

#### Experimental design

Corticosterone levels were manipulated orally using a minimally invasive technique described by Breuner et al. (1998), in which corticosterone dissolved in peanut oil was injected into mealworms *Tenebrio molitor*. We used a randomized block design with two doses of exogenous corticosterone and a control manipulation for each block.

The day before experimental day, we removed the food from the birds 1.5 hour before sunset and the mobile dividers were inserted into the cages to keep the birds individually. Water was still available *ad libitum*. The next morning (between 8:00 - 9:00, to avoid daily variation in hormone levels and to standardize the duration of the food removal), the experimenter quietly entered the room (the cages were oriented in a way that the birds could not see the door) and gave one mealworm to the selected bird through a small hole covered by a semi-transparent layer at the back of the cages, so that the bird could not see the experimenter, but we could observe the birds and record the time when they consumed the mealworm (mean time of ingestion was 39 seconds). We paid particular attention to avoid any visual or acoustic contacts with the birds. Fifteen minutes after the bird consumed the mealworm, the bird was captured through a backdoor at the cages and a single blood sample (~70μl) was taken as soon as possible. After blood sampling, body mass (to the nearest 0.1g) of the sampled individuals was also recorded and they were released back to their cage. The exact times when we entered the room, when the mealworm was given, when the bird ate the mealworm, when we caught the birds in the cages and when blood sampling was completed were all recorded. Handling time was defined as the time between when we opened the door of individual cages and blood sample collected. Total procedural time was defined as the time elapsed between the experimenter entering the room and when blood sampling was completed. Treatments were carried out in blocks, so that 3 birds in a block got the mealworms subsequently (with approximately 1 minutes staggering). Two blocks were sampled in a morning. After the treatments and sampling, the birds received the usual ad libitum bird chow and fresh water and were left undisturbed for the rest of the day. After we processed all birds, the experiment was repeated one week later, in which each individual received a different treatment than in the first trial, hence all birds received 2 different treatments. The treatments for the second trial were randomized again within blocks and birds were randomly assigned to the blocks.

#### Mealworm injection

Mealworms were injected with 20μl of peanut oil (VWR catalogue number: ACRO416855000) containing one of the following concentrations of corticosterone (Sigma catalogue number: NET399250UC): (1) control, no corticosterone; (2) low corticosterone, 0.2mg/ml (4μg corticosterone + 16μl peanut oil); and (3) 0.5mg/ml corticosterone concentration (10μg corticosterone + 10μl peanut oil). We made a stock solution for every concentration before we started the experiment. Hereafter we used those stock solution for injection. After thoroughly vortexing the solution, we injected it into mealworms with a 1ml syringe using a 26G-needle. The chosen corticosterone concentrations were based on previous studies and were calculated as dose per body mass. In red-eyed vireos *Vireo olivaceus* (12-16g), plasma corticosterone concentrations were elevated with a 0.2mg/ml concentration corticosterone solution (which corresponds to 0.28μg corticosterone per 1g of body mass) (Löhmus, Sundström & Moore, 2006). In Gambel’s white crowned sparrows *Zonotrichia leucophrys gambelii* (25-28g) a low (0.2mg/ml - 0.15μg/g) and a high (1mg/ml - 0.7μg/g) dose were used, but the low dose did not elevate significantly the plasma corticosterone levels (Breuner, Greenberg & Wingfield, 1998). Nestling Zebra-finches *Taeniopygia guttata* (6-15 g) received a dose of 0.25mg/ml (1.19μg/g) corticosterone, which resulted in a significant increase in circulating corticosterone levels (Spencer & Verhulst, 2008). Therefore, our manipulations correspond to a dose of 0.25μg/g (low) and 0.62μg/g (high).

### 3. Blood sampling

We took blood samples by puncturing the brachial vein with a 26G-needle and collecting blood in heparinized capillary tubes. Samples were kept on ice until transferring them to the lab (1-hours in the field and max. 1 h in the experiment). Samples were centrifuged at 2200g for 10 minutes and the plasma was removed with a Hamilton syringe. We divided the plasma into 2 aliquots, one for IGF-1 (15μl) and one for corticosterone (15μl). We stored the samples at −20°C until assayed for corticosterone by radioimmunoassay (RIA) and assayed for IGF-1 by enzyme-linked immunosorbent assay (ELISA).

### 4. Hormone assays

Plasma IGF-1 levels were measured in duplicates by a commercial avian ELISA kit (catalogue: cIGF1ELISA, lot number D00035) from IBT GmbH, Germany. The assay was developed to measure chicken IGF-1, and the amino acid sequence of IGF-1 is identical in chicken and in *P. biarmicus* (Á. Z. Lendvai et al. unpublished data), therefore this assay was expected to perform well in our study species. Serial dilutions of a plasma pool of *P. biarmicus* were parallel of the standard curve. IGF-1 was separated from its binding proteins using an acidic extraction in accordance with the manufacturer’s instructions. The final concentrations were determined colourimetrically measuring the absorbance at 450 nm using a Tecan F50 microplate reader. In this assay, we also included chicken *Gallus gallus* plasma samples as a reference, and the obtained IGF-1 concentrations for the chicken samples (375.2 ± 69.4 SE ng/ml) were an order of magnitude higher than what we had expected based on the literature (30-50 ng/ml, Ballard et al., 1990). Therefore we measured known concentrations of an international IGF-1 gold standard (WHO/NIBSC 02/254, a product used in different laboratories to calibrate IGF-1 values - Chanson et al. 2016) and recalibrated all concentrations against this standard. IGF-1 concentrations of the recalibrated chicken reference samples (53.0 gn/ml) were similar to published results and were in agreement with an in-house ELISA developed in our laboratory (Á. Z. Lendvai et al. unpublished data). Therefore, we used the recalibrated values in our analyses. Note however, that the recalibration only affects the absolute values reported, since the concentrations obtained originally would yield the same results, albeit on a different scale. Minimal detection limit was 1.5 ng/ml, and none of the samples fell below this limit. Intra-assay coefficient of variation was 3.9%, inter-assay CV was 5.7%.

Total corticosterone from plasma samples was quantified through direct radioimmunoassay (Lendvai, Bókony & Chastel, 2011). We extracted the corticosterone from plasma using diethylether, and extracts were reconstituted in phosphate-buffered saline. We let the samples incubate overnight at 4°C. We added ~10K dpm of 3H-Cort (Perkin Elmer Company: Catalogue number: NET399250UC, lot number: B00025), antiserum (Sigma C8784-100ST, lot number: 092M4784) and phosphate-buffered saline. We let the solution sit overnight at 4°C. Dextran-coated charcoal was added to separate corticosterone bound to antibodies. After centrifugation, the radioactivity of the bound fraction was counted in a liquid scintillation counter (QuantaSmart). All samples were processed in one assay (intra-assay CV: 3.5%, inter-assay CV: 5.4%).

### 5. Statistical analyses

We analysed our data in R statistical environment, R version 3.3.2 (R Core Team 2017). We fitted linear mixed models with function ‘lmer’ from package lme4 (Bates et al., 2014; version: 1.113), and we used stepwise backward model selection to find the best fitting model. Degrees of freedom for linear mixed models were calculated using the Satterthwaite approximation and corresponding p-values were obtained using the package lmerTest (Kuznetsova, Brockhoff & Christensen, 2016; version: 2.0-33).

To test the effects of handling stress on IGF-1 and corticosterone, we used generalized linear mixed models with individual as a random intercept. In the initial model, IGF-1 was the dependent variable, handling (first or second sample) and its two-way interaction with body mass and sex were the explanatory variables, and ring number (as individual identity) was the random factor. The same initial model structure was used to model corticosterone levels. We also analysed the relationship between IGF-1 levels and corticosterone levels in the first sample with a linear model controlling for body mass. Next, we calculated the stress-induced change in both corticosterone and IGF-1 levels (values from the first sample subtracted from the second samples), and analysed whether the magnitude of change between these two hormones were related in a linear model. In this analysis, the dependent variable was the change in IGF-1 levels and the explanatory variable was the change in corticosterone levels, while controlling for body mass and sex.

The experimental data were analysed using linear mixed models. Here, we considered treatment as a three-level factor (high corticosterone, low corticosterone and control). In the initial model, IGF-1 or corticosterone was the dependent variable, treatment and its two-way interaction with sex and body mass were the explanatory variables. The treatment blocks and the weeks of the experiment were also included as fixed effects. In both models we controlled for the handling time and total procedural time.

## Results

### Field samples

The 15 minute capture-restraint stress induced a significant increase in corticosterone levels (Fig.1a, Table 1), and body mass was positively related with corticosterone levels in the first sample (Table 1). Sex and the two-way interaction between stress and sex or body mass did not affect corticosterone levels (Table 1). Handling stress caused a significant decrease in IGF-1 levels (Fig.1b, Table 2), and IGF-1 levels were higher in males (Table 2). Body mass was not related to IGF-1 levels in the first sample. Body mass and the two way interaction between stress and sex or body mass did not affect the IGF-1 levels (Table 2).

**Table 1.** Parameter estimates of variables affecting circulating corticosterone levels in free-living bearded reedlings (*Panurus biarmicus*). Results are from the final linear mixed-effects model after stepwise backward elimination of non-significant effects. The initial model structure was: Corticosterone ~ Handling x (Sex + Mass). The terms excluded during model selection with the associated p-values in the model before elimination are shown below the table.

|  | Estimate | Std. Error | df | t-value | p-value |
| --- | --- | --- | --- | --- | --- |
| Intercept | -22.207 | 7.701 | 10.843 | -2.884 | 0.015 |
| Handling (stress) | 2.619 | 0.570 | 10.664 | 4.593 | <0.001 |
| Mass | 1.678 | 0.500 | 10.833 | 3.356 | 0.006 |

Terms excluded:
Handling\*Sex p=0.458, Sex p=0.401, Handling\*Mass p=0.096

**Table 2.** Parameter estimates of variables affecting circulating corticosterone levels in free-living bearded reedlings (*Panurus biarmicus*). Results are from the final linear mixed-effects model after stepwise backward elimination of non-significant effects. The initial model structure was: IGF-1 ~ Handling x (Sex + Mass). The terms excluded during model selection with the associated p-values in the model before elimination are shown below the table.

|  | Estimate | Std. Error | df | t-value | p-value |
| --- | --- | --- | --- | --- | --- |
| Intercept | 55.666 | 3.217 | 13.809 | 17.306 | <0.001 |
| Handling (stress) | -6.698 | 1.173 | 13.235 | -5.709 | <0.001 |
| Sex (males) | 14.181 | 3.880 | 13.001 | 3.655 | 0.003 |

Terms excluded:
Handling\*Mass p=0.644, Handling\*Sex p=0.507, Mass p=0.313

We did not find a significant relationship between IGF-1 and corticosterone levels in the first samples (t = 0.786, p = 0.450). The handling induced change in IGF-1 and in corticosterone (t = 0.669, p = 0.520) were also unrelated, even after controlling for body mass or sex (p’s > 0.9). One bird showed an unusual pattern in the corticosterone data, in which corticosterone decreased in response to handling stress. Removing this data point did not alter any of our conclusions.

**Figure 1:**
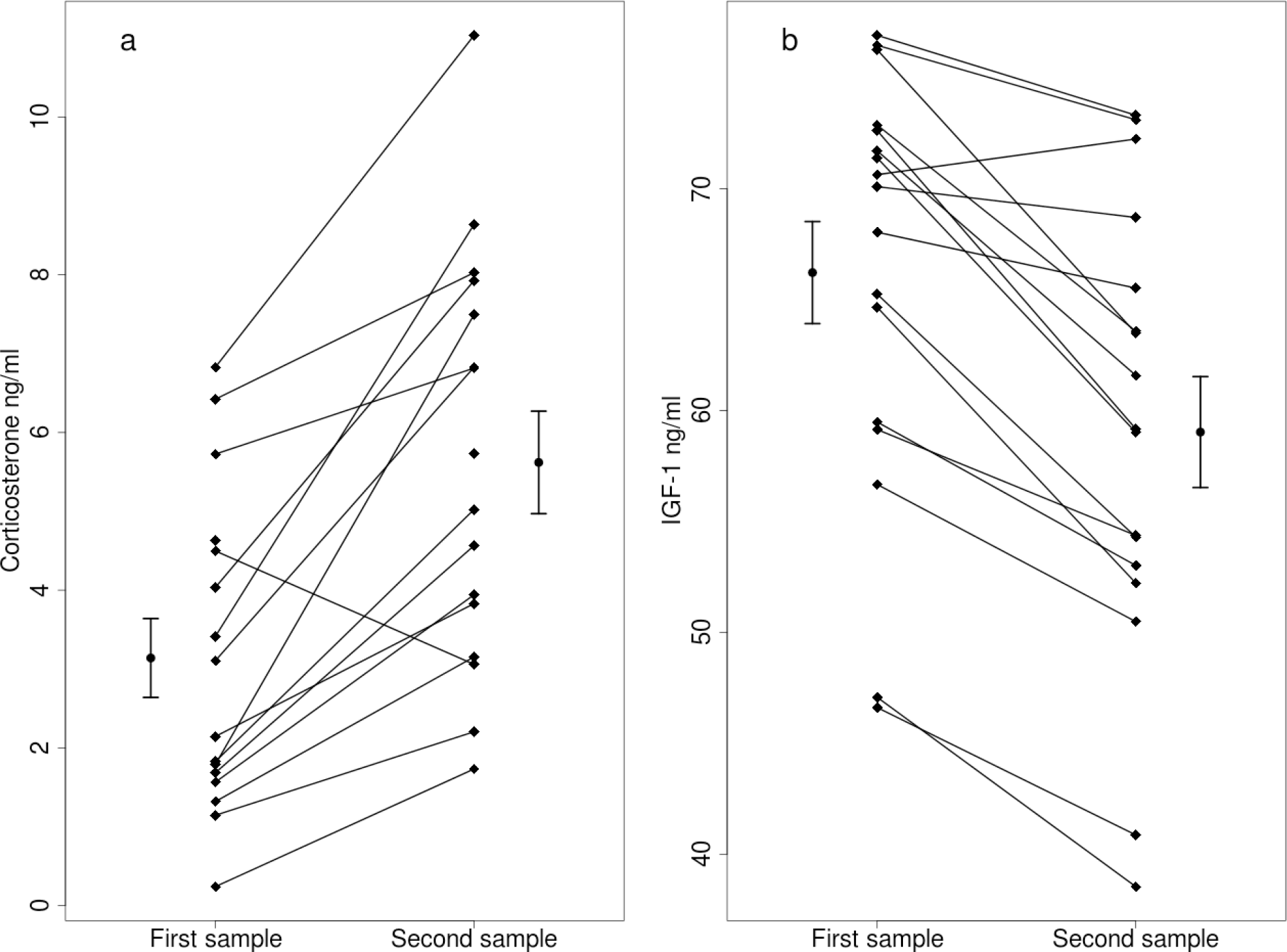
Capture handling stress causes (a) a significant increase in circulating corticosterone levels (n = 16) and (b) a significant decrease in circulating IGF-1 levels (n = 17) in free-living bearded reedlings (*Panurus biarmicus*). Squares denote the individual IGF-1 and corticosterone concentrations, and lines between the squares connect the first and second samples from the same individual. The dots and arrows beside the individual points represent the mean and standard error values of corticosterone (a) or IGF-1 (b) levels.

### Corticosterone manipulation

The body mass of the captive birds did not change during the two experimental weeks, and IGF-1 levels, corticosterone levels, block or sex were all unrelated to body mass (p’s > 0.5). The experimental week or the treatment blocks did not affect corticosterone or IGF-1 levels (p’s > 0.2).

Both low and high dose of treatment significantly increased corticosterone levels compared to the control treatment (Fig.2a, Table 3). Other effects, including sex, mass and the two-way interaction of treatment with sex and mass, handling time and total procedural time did not influence corticosterone levels (Table 3). Although the manipulation was successful in creating differences in circulating corticosterone levels, the treatment did not affect IGF-1 levels (Fig.2b, Table 4). Furthermore, sex, mass and the two-way interaction of treatment with sex and mass, handling time and total procedural time did not affect IGF-1 levels during the experiment (Table 4).

**Table 3.** Parameter estimates of variables affecting circulating corticosterone levels after oral administration of corticosterone in bearded reedlings (*Panurus biarmicus*). Results are from the final linear mixed-effects model after stepwise backward elimination of non-significant effects. The initial model structure was: Corticosterone ~ Handling time + Procedural time + Treatment x (Sex + Mass). The terms excluded during model selection with the associated p-values in the model before elimination are shown below the table.

|  | Estimate | Std. Error | df | t-value | p-value |
| --- | --- | --- | --- | --- | --- |
| Intercept | 5.660 | 4.550 | 28.750 | 1.244 | 0.223 |
| Treatment (low) | 18.260 | 6.100 | 33.790 | 2.993 | 0.005 |
| Treatment (high) | 43.100 | 6.100 | 33.790 | 7.006 | <0.001 |

Terms excluded:
Treatment (Low)\*Mass p=0.685, Treatment (High)\*Mass p=0.958,
Treatment (Low)\*Sex p=0.543, Treatment (High)\*Sex p=0.578, Sex p=0.530, Total procedural time p=0.267,
Handling time p=0.338, Mass p=0.249

**Table 4.** Parameter estimates of variables affecting circulating corticosterone levels after oral administration of corticosterone in bearded reedlings (*Panurus biarmicus*). Results are from the final linear mixed-effects model after stepwise backward elimination of non-significant effects. Treatment was part of the experimental design, so were kept in the final model, despite being not-significant. The initial model structure was: IGF-1 ~ Handling time + Procedural time + Treatment x (Sex + Mass). The terms excluded during model selection with the associated p-values in the model before elimination are shown below the table.

|  | Estimate | Std. Error | df | t-value | p-value |
| --- | --- | --- | --- | --- | --- |
| Intercept | 52.865 | 2.900 | 5.670 | 18.228 | <0.001 |
| Treatment (low) | -0.932 | 3.862 | 38.000 | -0.241 | 0.810 |
| Treatment (high) | 0.709 | 3.862 | 38.000 | 0.184 | 0.855 |

Terms excluded:
Handling time p=0.934, Treatment (Low)\*Mass p=0.911, Treatment (High)\*Mass p=0.729, Mass p=0.620,
Total procedural time p=0.502, Treatment (Low)\*Sex p=0.795, Treatment (High)\*Sex p=0.052, Sex p=0.561

## Discussion

We explored the mechanistic link between the HPA and HPS axes using a free-living songbird, resulting in two key findings. First, we found that in response to a standardized stressor, circulating IGF-1 levels decreased in wild bearded reedlings within 15 minutes. To our knowledge, this is the first study that reports such an effect for any free-living species. Second, we found that experimentally elevated corticosterone levels did not result in a decrease in IGF-1 levels during the same time frame as we found in the field. This result suggests that the somatotropic axis may respond to environmental stimuli independently from the HPA axis and may be part of the adaptive physiological coping mechanisms used to maintain or restore homeostasis in stressful situations.

**Figure 2:**
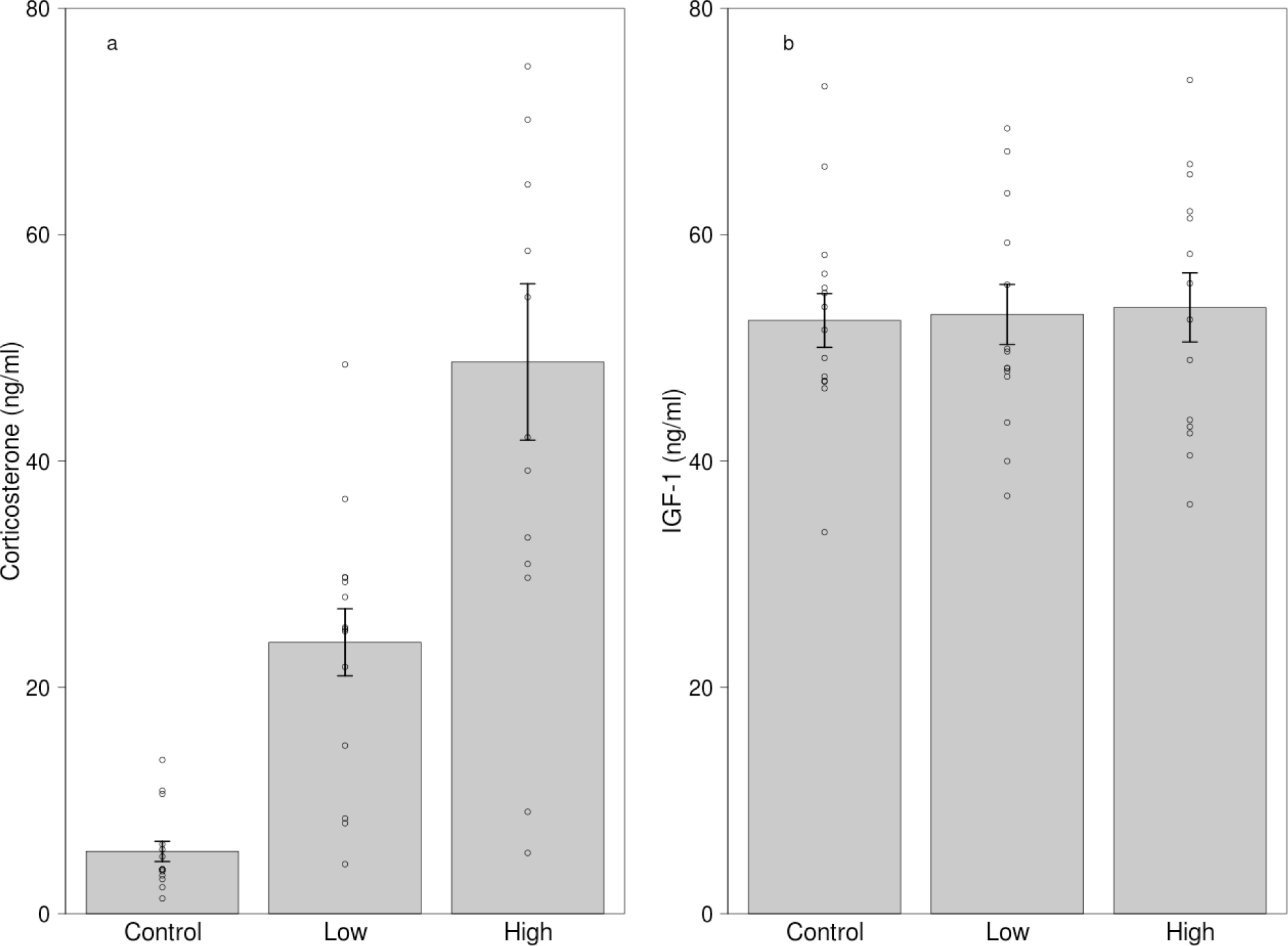
Effects of dietary corticosterone treatment on circulating (a) corticosterone (n = 42) and (b) IGF-1 levels (n = 42). Corticosterone levels were significantly higher in the low corticosterone (n = 14) and the high corticosterone (n=14) group compared with the control group (n = 14), although the IGF-1 levels did not differ between the treatment groups. We used 21 individuals in the experiment, and every bird received two different treatments.

Increases in plasma corticosterone levels in response to stress have been previously reported in many species. Our data obtained in bearded reedlings in the field is consistent with these finding, showing that corticosterone levels increase in response to capture/restraint stress and suggest that the birds perceived the procedure as stressful. Although many studies collect the stress-induced blood sample at 30 minutes (Wingfield, Vleck & Moore, 1992; Remage-Healey & Romero, 2001; Buehler et al., 2008), we chose to reduce the restraint period to study the effects of short-term acute stress on IGF-1 levels, while minimizing the potential confounding downstream effects of corticosterone. The effects of glucocorticoids are mainly genomic, which typically act over an hour, but require at least 15 minutes (reviewed in Haller, Mikics & Makara, 2008). Some short-term direct effects of corticosterone have been also demonstrated, such as effects on RNA synthesis (reviewed in Haller, Mikics & Makara, 2008). Therefore, by choosing a shorter restraint period, we aimed at decreasing the time during which the organism may have been exposed to the physiological effects of elevated glucocorticoids (Buehler et al., 2008). Although the effects of stress on IGF-1 levels in free-living organisms have not been reported before, this finding is consistent with previous studies in captive animals (Farmer et al., 1991; Wilkinson et al., 2006; Davis & Peterson, 2006; Wirthgen et al., 2017). However, our findings do not support the conclusion of earlier avian studies (Lodjak, Mägi & Tilgar, 2014; Lodjak, Tilgar & Mägi, 2016; Lodjak et al., 2017), in which handling time was reported to be unrelated to IGF-1 levels, although in nestlings. Such effects may be age-specific. For example, gilthead seabream *Sparus aurata* differed in expression of IGF-1 and IGF-1R mRNA levels during ontogeny (Perrot et al., 1999). In a study on Korean native ogol chicken, circulating IGF-1 levels gradually increased during the posthatching period (Yun et al., 2005). The stress-responsiveness of IGF-1 may also vary with development, and may explain why Lodjak, Mägi & Tilgar (2014) and Lodjak, Tilgar & Mägi (2016) did not find that IGF-1 was affected by handling. Furthermore, stress-responsiveness of IGF-1 may also be species-specific, although we also found that capture-restraint stress caused a decrease in IGF-1 levels in adult free-living house sparrows *Passer domesticus* (C. I. Vágási et al. unpublished data). Therefore, we suggest that further studies of IGF-1 levels should take into account the potentially confounding effect of additional stressors.

The decrease in IGF-1 under stress is consistent with the allostatic concept of the stress response (McEwen & Wingfield, 2003). Under stressful situations, the organism has to re-establish homeostasis and in order to do so, it has to suppress energetically costly anabolic processes and reinforce those behavioural and physiological processes that promote immediate survival (Wingfield et al., 1998). IGF-1 is a prime regulator of anabolic processes and antagonistic of the catabolic effects of glucocorticoids, therefore the decrease of IGF-1 under acute stress is consistent with its role as one of the physiological mechanisms responsible for maintaining homeostasis. For instance, in a previous study in mice, IGF-1 levels decreased markedly in food restricted animals and the individuals started to lose weight (O’sullivan et al., 1989). However, experimental IGF-1 administration during starvation reduced the rate of weight loss through the inhibition of the catabolic processes (O’sullivan et al., 1989). The sudden drop of IGF-1 levels in response to the stressor suggests that this physiological change prepares the animal for the metabolic challenges faced by slowing down anabolic processes and permitting the catabolic effects of glucocorticoids.

In light of these results, we expected that higher glucocorticoid stress responses would be associated with the largest decrease in IGF-1 levels. However, despite the opposite direction of change in the two hormones, neither levels in the first sample nor the magnitude of corticosterone and IGF-1 stress responses were related in the field study at the individual level: birds with the strongest corticosterone increase were not the ones that decreased their IGF-1 levels the most, and vice versa. Furthermore, we did not find any relationship between those parameters if we controlled for body mass. In order to test the relationship between the two hormones more thoroughly, we carried out an experiment in which we manipulated corticosterone in a minimally invasive manner.

Our dietary hormone treatment increased the circulating levels of corticosterone, while control birds did not show a marked increase in corticosterone levels over the course of the study. These results suggest that similarly to previous studies (Breuner, Greenberg & Wingfield, 1998; Löhmus, Sundström & Moore, 2006; Spencer & Verhulst, 2008), our oral hormone treatment was successful. Corticosterone concentrations after ingesting the mealworm were significantly higher in both the low and the high dose group compared to the controls, albeit with large individual variation. Despite this rapid increase in corticosterone levels in the absence of a physical stressor, IGF-1 levels in our treated birds remained at the level of the controls, with minimal effect sizes; therefore, we can be confident that the treatment did not affect IGF-1 secretion. Our results are similar to those reported in the sunshine bass, in which confinement stress for 15 minutes resulted in a decrease in IGF-1 levels, but the dietary hormone treatment did not affect plasma IGF-1 concentrations (Davis & Peterson, 2006).

These results suggest that the HPA and HPS axes are not linked downstream at the endpoint of these hormonal cascades, but the crosstalk between these pathways happens at the hypothalamic-pituitary level. We argue that this is the reason behind the discrepancy between the conclusions of studies using physiological and pharmacological doses (see above). Dexamethasone, a powerful glucocorticoid agonist, which is known for having a strong negative feedback on the HPA axis has also been shown to decrease plasma IGF-1 concentration in chickens (Leili & Scanes, 1998), which supports the notion that integration of the HPA and HPS axes is operating at higher regulatory levels.

According to a recent study performed on great tit nestlings, the relationship between corticosterone and IGF-1 varies with the nutritional condition of the individuals. Lodjak, Tilgar and Mägi (2016) found that pre-fledging plasma IGF-1 levels of nestlings in good condition (from broods that were experimentally reduced) were positively related to feather corticosterone (an integrated measure of corticosterone over several days during the development), whereas the association between IGF-1 and feather corticosterone levels was negative in nestlings with lower nutritional condition (from enlarged broods). In control broods however, there were no association between the two hormones. In our captive study, the birds had *ad libitum* food availability and they were all in good condition; therefore, one possible explanation for the absence of the relationship between IGF-1 and corticosterone may be that birds in good condition can afford to tolerate higher glucocorticoid concentrations without decreasing IGF-1 levels. In line with this possibility, biomedical studies have shown that IGF-1 diminishes the protein catabolic effects of glucocorticoids only in normally fed but not in starved subjects (Botfield, Ross & Hinds, 1997).

## Conclusions

In this study, we showed that IGF-1 levels decrease in response to stress in a free-living songbird, and that the magnitude of this response is not related to the glucocorticoid stress response. Furthermore, an experimental increase in corticosterone did not affect circulating IGF-1 levels. While glucocorticoids may still have non-linear or permissive effects on IGF-1 regulation, our results suggest that the HPA and HPS axes are both stress responsive and are not tightly coregulated at their downstream endpoints. These results raise the possibility that the interaction between IGF-1 and corticosterone may modulate the adaptive response of organisms in stressful situations. Investigations of the relationship between glucocorticoids and fitness remain equivocal, with some studies showing positive, negative and also no relationship (reviewed in Bonier et al., 2009). The lack of a general glucocorticoid-fitness relationship has been suggested to be a result of the flexibility and environmentally context dependent nature of glucocorticoids (Bonier & Martin, 2016). Our results showing that IGF-1 levels are responsive to stress independently from glucocorticoids suggest that the HPS axis is an autonomous physiological cascade that may be also involved in the mediation of life history decisions and affect fitness components (Harshman & Zera, 2007; Dantzer & Swanson, 2012; Lewin et al., 2016). IGF-1 is an evolutionary ancient regulatory hormone with a primary role to provide an organism-wide internal signal about resource availability, and may alter the function of glucocorticoids. If overall resource availability is high (as was the case in our captive study), then IGF-1 levels can act as a physiological buffer against the adverse effects of increased corticosterone levels. However, when the central nervous system receives input from the environment challenging the organism, it may require the reallocation of resources, which is reflected in decreased IGF-1 levels. Therefore, circulating IGF-1 levels can be an important biological indicator of individual internal state and a useful parameter to investigate in the study of life-history decisions.

## Acknowledgements

We thank Gyula Ölveczki, Janka Pénzes and Anna Anita Rácz for excellent help in the fieldwork and the experiment. Scott MacDougall-Shackleton and an anonymous reviewer provided constructive comments on an earlier version of the manuscript. We are grateful for the Hortobágy National Park to make the work possible.

